# aRNApipe: A balanced, efficient and distributed pipeline for processing RNA-seq data in high performance computing environments

**DOI:** 10.1101/060277

**Authors:** Arnald Alonso, Brittany N. Lasseigne, Kelly Williams, Josh Nielsen, Ryne C. Ramaker, Andrew A. Hardigan, Bobbi Johnston, Brian S. Roberts, Sara J. Cooper, Sara Marsal, Richard M. Myers

**Affiliations:** HudsonAlpha Institute for Biotechnology, Huntsville, AL 35806, USA; Rheumatology Research Group, Vall d’Hebron Hospital Research Institute, Barcelona, Spain; Department of Genetics, The University of Alabama at Birmingham, Birmingham, AL 35294, USA.

## Abstract

**Summary:** The wide range of RNA-seq applications and their high computational needs require the development of pipelines orchestrating the entire workflow and optimizing usage of available computational resources. We present aRNApipe, a project-oriented pipeline for processing of RNA-seq data in high performance cluster environments. aRNApipe is highly modular and can be easily migrated to any high performance computing (HPC) environment. The current applications included in aRNApipe combine the essential RNA-seq primary analyses, including quality control metrics, transcript alignment, count generation, transcript fusion identification, alternative splicing, and sequence variant calling. aRNApipe is project-oriented and dynamic so users can easily update analyses to include or exclude samples or enable additional processing modules. Workflow parameters are easily set using a single configuration file that provides centralized tracking of all analytical processes. Finally, aRNApipe incorporates interactive web reports for sample tracking and a tool for managing the genome assemblies available to perform an analysis.

**Availability and documentation:** https://github.com/HudsonAlpha/aRNAPipe; DOI:10.5281/zenodo.202950

**Supplementary information:** Supplementary data are available at Bioinformatics online.

## 1 Introduction

Quantification of RNA transcripts by next-generation sequencing technologies continues to increase in both throughput and capabilities as sequencing becomes more affordable and accessible (McGettigan, 2013). Unlike gene expression microarrays, RNA-seq not only quantifies gene expression levels, but also measures alternative splicing, transcript fusions, and RNA sequence variants (Finotello and Di Camillo, 2015; Koboldt, et al., 2012; Maher, et al., 2009). This broad spectrum of applications has fostered development of a rich set of bioinformatics methods focused on each processing stage (Conesa, et al., 2016). Current RNA-seq data primary analysis applications usually apply a single processing step, involve complex dependencies between processing stages, and depend on the sequencing protocol performed (see Supplementary Section 1). Consequently, there is an increasing need for tools orchestrating the analysis workflow to ensure repeatability of RNA-seq data processing. In addition to the need for data processing integration, the computational requirements of some RNA-seq analysis steps are a bottleneck (Scholz, et al., 2012) and, the use of high performance computing (HPC) clusters is unavoidable. Because HPC clusters are a valuable and often limited resource, tools integrating RNA-seq processing stages must be carefully designed and optimized. Considering these challenges, we developed a balanced, efficient and distributed pipeline for RNA-seq data analysis: aRNApipe (**a**utomated **RNA**-seq **pipe**line). This pipeline was optimized to efficiently exploit HPC clusters, to scale from tens to thousands of RNA-seq libraries, and includes modules yielding complete RNA-seq primary analysis.

## 2 Methods

aRNApipe is designed to overcome the challenges of integration, synchronization and reporting of RNA-seq data analysis by using a project-oriented and balanced design optimized for HPC clusters (Fig. 1).

The core application of aRNApipe (Supplementary Section S1) includes six operating modes: 1) executing a new analysis, 2) updating a previous analysis to include new samples or enable new modules, 3) showing analysis progress, 4) building a project skeleton, 5) showing available genome builds, and 6) stopping an ongoing analysis.

### Input data

aRNApipe requires two input files: 1) analysis configuration, and 2) samples to include in the analysis. In the configuration file, the user can set the executing parameters, including enabling/designating arguments of processing modules, assigning computational resources to modules, and selecting the reference genome build.

### Current applications

aRNApipe currently includes applications covering the main variations of RNA-seq data generation (Supplementary Section S2). Throughout the workflow, a main daemon process manages pipeline execution (i.e. inter-dependencies between applications) and monitors analysis of each sample at each stage (Fig. 1). First, low-quality reads/bases and adapter sequences can be filtered. Then, a second stack of applications is run in parallel, including assessment of raw data quality, transcript alignment and quantification and identification of gene fusions. The main process launches a third stack of analyses including quantification of genes and exons, conversion of SAM files to BAM sorted files, and alignment quality. Finally, a fourth stack of applications including variant calling and alternative splicing modules are run.

**Fig. 1.**
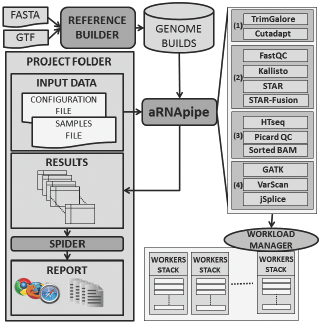
aRNApipe workflow for primary analysis of RNA-seq data

### Report generation

The Spider is an aRNApipe add-on module that generates interactive web reports summarizing an aRNApipe analysis. These reports review each module, including sample quality control metrics (Supplementary Figures 1 and 2). The user can also observe the computational resources used by each module and access all logs generated during analysis (Supplementary Figure 3). Additionally, the Spider generates matrix-like count data files of raw counts, RPKMs, and corresponding annotation files (i.e. gene identifier and length). Supplementary Section S3 provides a list of generated outputs.

### Reference builder

The programs used for RNA-seq data processing often use different formats and standards. To address this problem and to provide a centralized repository of available genome builds, aRNApipe includes a reference builder that generates all required files for a genome build based on initial files obtained from sources like NCBI and Ensembl repositories (Supplementary Section S4 and Supplementary Figure 4).

### Implementation

aRNApipe has been developed using Python 2.7 (Supplementary Figure 5). When running on an HPC cluster, aRNApipe relies on the workload management application to submit jobs for each processing stage, taking into account cross-stage dependencies and custom resource requirements for each stage. aRNApipe has been implemented with the workload management system IBM Platform LSF, but its design allows quick migration to any other workload manager by editing one Python library (Supplementary Section S5). Additionally, a single-machine version is also provided. A configuration library provides supply paths to all applications used by aRNApipe.

## 3 Results

We have extensively tested aRNApipe and used it to analyze hundreds of RNA-seq libraries with multiple configurations, including different species, different genome builds and different RNA-seq protocols. The reports generated for four example datasets can be accessed online (http://arnapipe.bitbucket.org): (1) Strand-specific paired-end RNA-seq data from 9 human samples of different tissues (GSE69241), (2) unstranded single-end data from two paired normal and colorectal tumor tissues (GSE29580), (3) unstranded paired-end data from 11 melanoma samples (GSE20156) and 1 prostate cancer cell line (NCIH660), and (4) in-house unstranded paired-end data from 20 zebrafish libraries.

## 4 Conclusions

aRNApipe provides an integrated and efficient workflow for analyzing single-end and stranded or unstranded paired-end RNA-seq data. Unlike previous pipelines, aRNApipe is focused on HPC environments and the independent designation of computational resources at each stage allow optimization of HPC resources (see Supplementary Section 1). This application is highly flexible because its project configuration and management options. The Spider provides functional reports for the user at all analytical stages and the reference builder is a valuable genome build manager. Implementation of this pipeline allows users to quickly and efficiently complete primary RNA-seq analysis.

## Acknowledgements

We thank the Myers Lab at the HudsonAlpha Institute of Biotechnology for their constructive suggestions, feedback and testing during implementation of this pipeline. We also thank the HudsonAlpha IT team for their continuous support during the development of the pipeline and its optimization on the HPC cluster.

### Conflict of Interest

none declared.

### Funding

This work was supported by funds from The HudsonAlpha Institute for Biotechnology. Andrew Hardigan and Ryne Ramaker are funded by the NIH-National Institute of General Medical Sciences Medical Scientist Training Program (5T32GM008361-21). Brittany Lasseigne is funded by the William J. Maier III Fellowship in Cancer Prevention (Prevent Cancer Foundation).

